# Is laterality adaptive? Pitfalls in disentangling the laterality–performance relationship

**DOI:** 10.1101/694968

**Authors:** J. Paszulewicz, P. Wolski, M. Gajdek

## Abstract

Unlike non-human animal studies that have progressively demonstrated the advantages of being asymmetrical at an individual, group and population level, human studies show a quite inconsistent picture. Specifically, it is hardly clear if and how the strength of lateralization that an individual is equipped with relates to their cognitive performance. While some of these inconsistencies can be attributed to procedural and conceptual differences, the issue is aggravated by the fact that the intrinsic mathematical interdependence of the measures of laterality and performance produces spurious correlations that can be mistaken for evidence of an adaptive advantage of asymmetry. Leask and Crow [Leask, S. J., & Crow, T. J. (1997), How far does the brain lateralize?: an unbiased method for determining the optimum degree of hemispheric specialization. *Neuropsychologia*, *35*(10), 1381–1387] devised a method of overcoming this problem that has been subsequently used in several large-sample studies investigating the asymmetry–performance relationship. In our paper we show that the original Leask and Crow method and its later variants fall victim to inherent nonlinear dependencies and produce artifacts. By applying the Leask and Crow method to random data and with mathematical analysis, we demonstrate that what has been believed to describe the true asymmetry–performance relation in fact only reflects the idiosyncrasies of the method itself. We think that the approach taken by Leask in his later paper [Leask, S. (2003), Principal curve analysis avoids assumptions of dependence between measures of hand skill. *Laterality*, *8*(4), 307–316. doi:10.1080/13576500342000004] might be preferable.

*No part of the study procedures or analyses was pre-registered in an institutional registry prior to the research being conducted

## 1. Introduction

### 1.1 Is laterality adaptive?

The 19th-century discovery of the neural correlate of “aphemia” in the left posterior inferior frontal gyrus established a link between brain asymmetry and language (cf. Harrington, 1995). This notion was reaffirmed by ample empirical evidence including, but not restricted to, observations of anomalous laterality in language disorders (Orton, 1937; Travis, 1931). Gradually, the belief that brain asymmetry is a uniquely human prerequisite for language and is overtly manifested by the ubiquitous right-hand preference became paradigmatic. Despite some speculation on how brain asymmetry was adaptive (e.g. Levy, 1977), the question of *if* it was adaptive tended to be neglected. This changed at the turn of the 21st century due to the gradual erosion of the traditional understanding of laterality, and a move towards – as Corballis postulated – “a more realistic and biologically based understanding of what cerebral asymmetry is really about” (1998, p. 152). Most of the studies (reviewed in Rogers, 2014; Ocklenburg & Güntürkün, 2012) and theorizing (Ghirlanda & Vallortigara, 2004; Rogers, Vallortigara, & Andrew, 2013) on the evolution and adaptive advantages of laterality have been done with non-human animals, but there have been human studies, too. Naturally, these studies concentrate more on the advantages or potential evolutionary adaptiveness of laterality than on its evolution. In one of the first such studies, Boles and colleagues (Boles and Barth, 1999; 2008) explored the relationship between individual differences in the degree of asymmetry and the level of performance. They tested a relatively large sample of subjects with a variety of cognitive tasks and found a number of positive correlations between the absolute values of asymmetry and performance. Surprisingly, of 13 tasks employed, for 8 the correlations were negative, precluding the generalization that performance increases with lateralization. Other studies diverge too: some report a positive relation (Crow, Crow, Done, & Leask, 1998; Mellet et al. 2014; Hirnstein, Hausmann, & Hugdahl, 2014), some a negative relation (Ladavas & Umilta, 1983; Hirnstein, Hausmann, & Gunturkun, 2008; Hausmann et al., 2019), and some no relation between asymmetry and performance at all (Lust, Geuze, Groothuis, & Bouma, 2011; Lust et al., 2011). Chiarello, Welcome, Halderman and Leonard (2009) and Chiarello, Vazquez, Felton, and Leonard (2013) found a positive relation between asymmetry and performance in lexical processing, but only in individuals with consistent hand preference. Crow et al. (1998) presented results suggesting that equal hand skill is detrimental to performance, while Hirnstein, Leask, Rose and Hausmann (2010) demonstrated the opposite tendency, i.e. several cognitive tasks were performed relatively better by subjects showing no or only mild asymmetry. Also, in mathematically gifted subjects symmetry has been found to be optimal for executive function, fluid reasoning (Desco et al., 2011), and mental rotation (O’Boyle et al., 2005). In the structural domain, Catani et al. (2007) found symmetry of *arcuate fasciculi* to be advantageous for verbal recall.

### 1.2 Spurious correlations

While the contradicting findings certainly demand an explanation, their interpretation is even harder due to an alarming possibility that many of the published laterality–performance correlations may be spurious. Leask and Crow (1997) discussed the consequences of the fact that the typically used measures of asymmetry in the form of either absolute *R* − *L* or relative (*R* − *L*)/(*R* + *L*) lateral differences are not mathematically independent of performance. In a similar vein, Mazoyer and Tzourio-Mazoyer (2004) demonstrated that the significant correlation between the right *planum temporale* surface area and the index of asymmetry of the temporal plana surfaces, as was observed in the classic study of Geshwind and Levitsky (1968; Galaburda, Corsiglia, Rosen, & Sherman, 1987), was a mathematical artifact due to the interdependence of both measures; they even provided mathematical formulae for predicting this and other similar spurious correlations. Although the authors were concerned with the part–sum relations of surfaces, their reasoning applies to all cases in which a meaningful relation is inferred from linear correlation of mathematically interdependent measures. These studies demonstrate clearly that attempts at establishing a relation between laterality and performance can – and indeed sometimes do – go astray due to the properties of the measures used as the operationalizations of these constructs.

### 1.3 The Leask and Crow method

In their important 1997 paper, Leask and Crow discussed the problems related to the analysis of experimental results caused by the fact that any causal influence of laterality on performance resulting from the purported advantage of asymmetric organization might be mixed with an inherent mathematical interdependence of the employed measures (Leask and Crow, 1997; 2006). To separate the effect of the true causal influence from the spurious dependence reflecting just the mathematical interdependence of the measures, they invented a method of data analysis (modified slightly in Leask & Crow, 2006) which they and their colleagues used in several studies (Leask & Crow, 1997; 2006; Hirnstein et al., 2010; Hirnstein et al., 2014). We refer to this algorithm as the “Leask and Crow method” or the LCM for short.

In order to filter out the inherent mathematical interdependence between Laterality Index *Z* = (*R* − *L*)/(*R* + *L*) and the measure of performance (*R* or *L*) (Leask & Crow, 1997, 2006), they first determine – with locally weighted scatterplot smoothing (LOESS) – local means of observed performance levels for different Laterality Index values, and then subtract similarly computed means for reference data. As a reference they use either randomly generated, uncorrelated data (Leask and Crow, 1997), or actual observations, decorrelated by displacing *R* and *L* values relative to each other (Leask and Crow, 2006). They believe that the resulting difference curve shows the net dependence of performance (measured by *R* or *L*) on asymmetry (expressed by the Laterality Index *Z*), free of the mathematical interdependence masking the true effect.

An example of application of the LCM to a large dataset of hand performance on a box-marking task (NCDS data^2^, same as used by Leask and Crow 1997; 2006) is presented in Figure 1A. The chart on the left shows the scatterplot of right-hand performance vs Laterality Index, *Z* (grey dots), with superimposed LOESS curves for the source (solid line) and randomly generated reference data (dashed line). The chart on the right presents the difference curve representing the alleged net dependence of performance on asymmetry, devoid of the effects of the mathematical interdependence. The curve displays a performance maximum for the Laterality Index value of ca 0.1, corresponding to mild right-handedness, which the authors consequently interpret as an “optimum degree of hemispheric specialization” (Leask & Crow, 1997).

**Figure 1.**
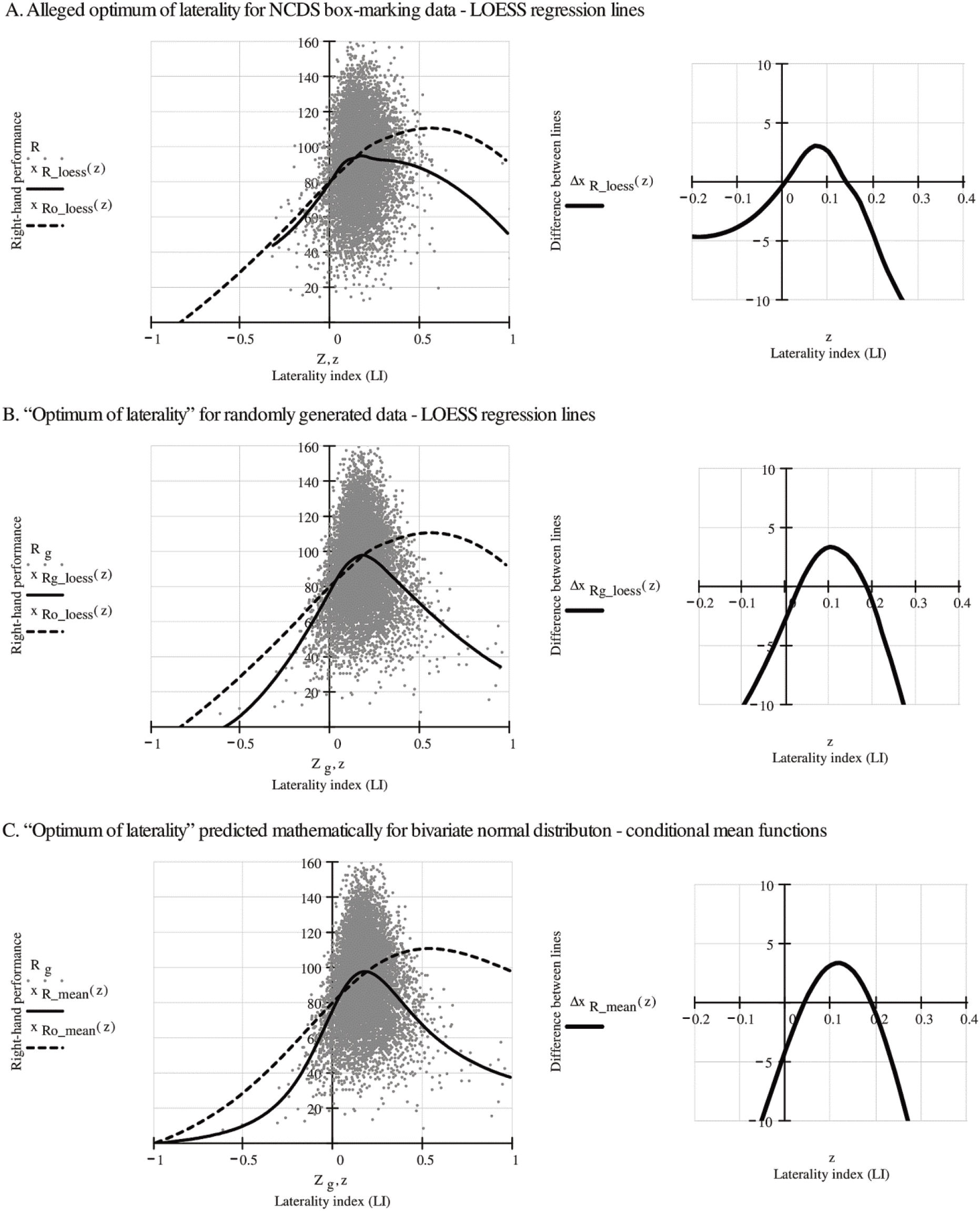
Leask and Crow method applied to: A) NCDS box-marking scores; B) randomly generated data; and C) analytically derived conditional mean functions. Grey scatterplots depict right-hand performance *R* against the lateralization index *Z* for the actual box-marking scores in chart A, and for random correlated dataset *R*_g_,*Z*_g_ with the same distribution parameters and correlation in charts B and C. The graphs on the left show: A) Local regression LOESS curves plotted for the actual box-marking scores; B) LOESS curves for simulated random data; and C) conditional mean functions for binormal distribution. Solid lines in these graphs show the curves *x*_R_loess_(*Z*), *x*_R_mean_(*Z*) for correlated (*ρ* = 0.782) data and the dashed lines show the curves *x*_Ro_loess_(*Z*), *x*_Ro_mean_(*Z*) for uncorrelated (*ρ* = 0) reference. The graphs on the right show difference functions Δ*x*_R_loess_(*Z*), Δ*x*_Rg_loess_(*Z*) and Δ*x*_R_mean_(*Z*) obtained by subtracting the dashed-line from the solid-line curves on the left. The similarity of the difference curves suggests that the allegedly unbiased depiction of the relation between laterality and performance provided by the LCM is mostly an artifact. Regression lines in graphs A and B were calculated with the built in LOESS regression procedure (span = 0.6) of Mathcad Professional. The values of conditional mean functions presented in graph C were calculated from Equations 15 & 18.

## 2. The problematic validity of Leask and Crow’s method

We believe that the picture of the alleged influence of laterality on performance produced by the LCM, including the observed “optimum of laterality” at mild right-handedness, is wrong. The mathematical dependence between the measure of performance (here *R*) and Laterality Index *Z* is nonlinear and differs for the zero and non-zero correlation between *R* and *L*. Hence, the difference curve (obtained by subtracting the local regression approximation computed for uncorrelated reference data from the local regression approximation computed for correlated real data) presents a mathematical artifact of the *R* and *L* correlation.

### 2.1 The LCM applied to box-marking data*

To demonstrate the problem with the LCM, we applied it to randomly generated test “performance scores”^3^ with distributions and correlation approximating real-life NCDS box-marking data. The results are presented in Figure 1B. Randomly generated “scores” obviously cannot reflect any real-life influence of laterality on performance. Therefore, for random data, the LCM should produce a picture quite different from the one obtained for real observations (Figure 1A); however, this was not the case. The charts in Figures 1A and B are strikingly similar: the run of the LOESS difference curve computed for the randomly generated “scores” very closely resembles what Leask and Crow obtained with NCDS data, including the effect they misinterpreted as manifestation of an optimum degree of laterality.

The presence of the extremum in question can be predicted analytically. The local regression used by the LCM produces lines that almost perfectly follow the course of conditional mean functions (cf. Figure 5 in the Appendix). As we know the underlying probability density functions (Equation 1) for variables *R*_g_ and *L*_g_ that simulate the performance “scores” of the two hands, we can examine the behavior of the LCM using analytically derived conditional mean functions to compute estimates of local expected values of right-hand performance *R* for fixed values of the lateralization index *Z*. Details of deriving the formulae (Equations 15 and 17) for conditional mean functions (and also for conditional modes) are provided in the appendix. The conditional mean functions for correlated (solid line) and uncorrelated (dashed line) reference data have been plotted, along with the scatterplot of the simulated right-hand performance data, in Figure 1C left. As in Figures 1 A and B, the right chart shows the difference curve.

A qualitative comparison of the three charts shows that the maximum of the difference function Δ*x*_R_loess_(*Z*) computed for NCDS box-marking data (Figure 1A right) is very similar (even if slightly noisier) to the maxima of difference function Δ*x*_Rg_loess_(*Z*) in randomly generated data (Figure 1B right) and the mathematically derived function Δ*x*_R_mean_(*Z*) (Figure 1C right)^4^. Because the latter two by definition involve no alleged real-life causal influence of asymmetry on lateralization, it follows that the first one should also be considered an artifact produced by the LCM algorithm rather than the true effect of causal influence of asymmetry on performance. The artifactual effect is mostly produced by the interaction of two factors: the choice of a nonlinear function (i.e. laterality index) of the original left- and right-hand performance data as an independent variable, and the adoption of uncorrelated data as a reference dataset. The interpretation of the maximum as an optimum of laterality does not therefore seem appropriate. The extrema of the regression lines computed for the true observations, and for the uncorrelated reference, as well as the maximum that results from subtracting the values of the two, should not be considered as much more than just mathematical artifacts. Even if some true causal effect of asymmetry were indeed hidden in the data, it would evidently have to be very small to be almost entirely masked by the LCM-generated artifact we see in Figure 1.

To sum up, comparing the effects of applying the LCM to real-life and random test data, we can conclude that the Leask and Crow method fails to remove the inherent mathematical interdependence of the measures of laterality and performance. Even worse, by subtracting two complex but different nonlinearities of the laterality index, the algorithm produces yet another nonlinear dependence which quite realistically mimics the effect of a possible causal influence of laterality on performance, with an apparent optimum at moderate rightward asymmetry.

### 2.2 The LCM applied to dichotic listening results

Hirnstein et al. (2014) published a paper reporting an application of the LCM method to a large set of lateralized dichotic listening performance measurements that was very different than the box-marking hand performance data analyzed by Leask and Crow (1997). They obtained a picture of the alleged relation between laterality and performance that was quite contrary to the one described by Leask and Crow. Instead of a single optimum, they found a u-shaped relation that was suggestive of an advantage in performance proportional to the absolute size of the asymmetry.

There are two critical differences between the data of Leask and Crow and the data of Hirnstein et al. In contrast to box-marking performance, which shows strong positive correlation between the hands, the left- and right-ear scores in a dichotic listening task correlate negatively; they are also expressed by percentages of correct responses that for each subject sum up to 100%. Decorrelation of such data – a part of the LCM algorithm, here performed by pairing left- and right-ear scores from different subjects – generates a nontrivial proportion of observation pairs with sums greater than 100% (cf. scatterplot in Figure 3B). This departure from the original task’s restrictions produces an unwarranted difference in distributions between the proper observations and the reference dataset. This discrepancy affects the final difference curve and adds to the problems already present in the LCM that we demonstrated earlier and elaborate on next.

As we show in Figure 2, the course of the difference curve produced by the LCM, representing the mathematical interdependence between laterality and performance measures (here derived from conditional mean functions), depends strongly on the direction and strength of the correlation between *R* and *L* performance scores ^5^. While positive correlation produces an artifactual “optimum” as observed by Leask and Crow (1997), for negative values of the correlation coefficient, an equally superficial effect that is disturbingly similar to the one found by Hirnstein et al. (2014) is present.

**Figure 2.**
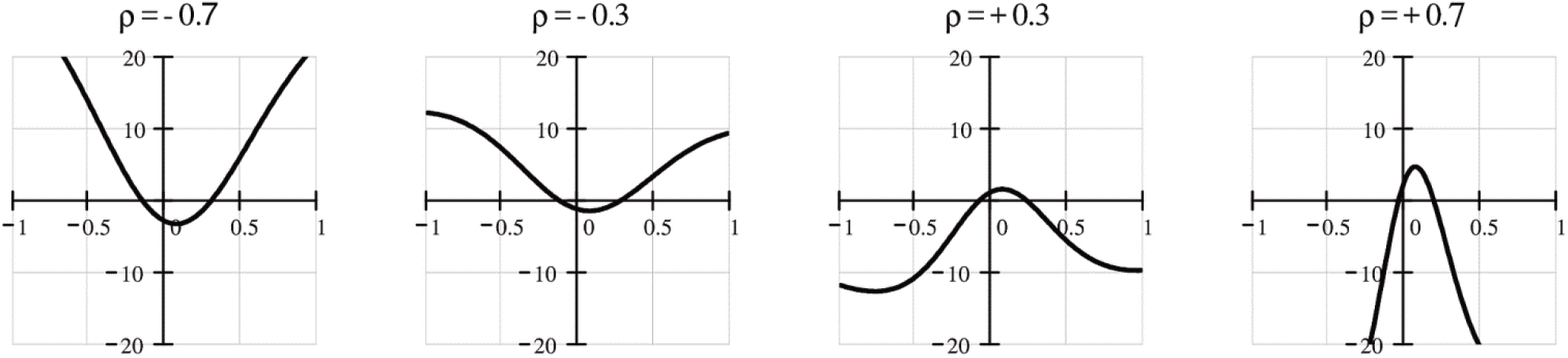
The effect of *R*, *L* correlation (*ρ*) size and direction on the course of the difference function Δ*S*_mean_(*Z*; *ρ*) = *S*_mean_(*Z*; *ρ*) − *S*_mean_(*Z*; 0) (vertical axes) versus the lateralization index *Z* (horizontal axes). The symbol *S*_mean_(*Z*; *ρ*) stands for the conditional mean of overall performance derived for the binormal distribution function of *R*, *L* parameterized by correlation *ρ* (cf. Equation 14).

Just as with the data of Leask and Crow, we tested the LCM algorithm used in Hirnstein et al. (2014) with randomly generated data that match the *R*, *L* correlation and distribution parameters of the original data.

The result of the simulation is presented in Figure 3A as a scatterplot of overall percent performance level *S* against lateralization index *Z*. Conditional mean function for overall performance 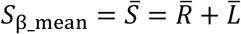 is plotted with a thin dashed line, while the local regression LOESS curve *S*_loess_(*Z*) is plotted with a solid line. The course of both functions is very similar to that of the local regression line for real data in the paper of Hirnstein et al. (2014, Fig. 1A). Following the authors’ procedure, we decorrelated the simulated data by shifting the *R* and *L* scores with respect to each other (see Appendix 4.5 for details). The obtained reference set of performance *S*_ref_ and laterality index *Z*_ref_ pairs is presented as a scatterplot in Figure 3B. As before, we also computed the conditional mean *S*_ref_mean_(*Z*) (dashed line), and local regression *S*_ref_loess_(*Z*) (solid line) functions. Figure 3C presents the differences of local regression curves *S*_loess_(*Z*) − *S*_ref_loess_(*Z*) (solid line) and mathematically derived conditional mean functions *S*_β_mean_ − *S*_ref_mean_(*Z*) (dashed line). Both show a relation very similar to the one obtained by Hirnstein et al. (2014, Fig 1D) with a minimum at LI values corresponding to mild rightward asymmetry. Being derived from simulated data, the obtained difference curve by definition represents nothing more than just the mathematically predictable consequences of applying the LCM to negatively correlated random observations. The similarity of the curves we obtained for simulated random data to those computed by Hirnstein et al. for true measurements persuasively shows that once again the LCM fails not only to remove the mathematical interdependence, but also to reveal the true causal influence of asymmetry on performance. Therefore, as in the case of the box-marking results, the interpretation of the obtained dependencies as true phenomena pertinent to the problem of adaptive value of laterality is misguided.

**Figure 3.**
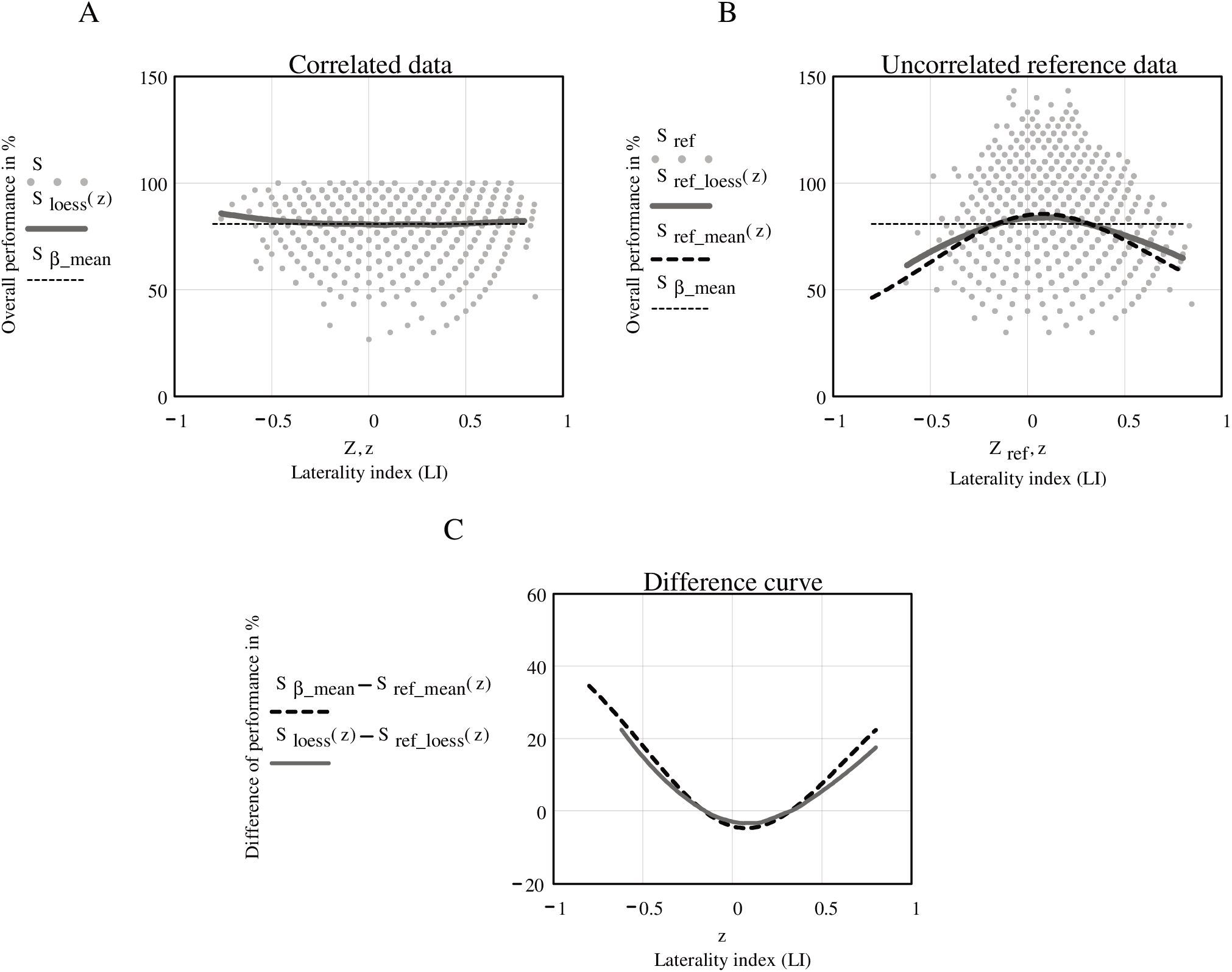
The Leask and Crow method applied to randomly generated data matching the parameters of dichotic listening results (non-forced condition) of Hirnstein et al (2014). A. Scatterplot of overall performance *S* against lateralization index *Z* for correlated data (Corr(*R*, *L*) = *ρ* ≅ −0.51), with LOESS regression line *S*_loess_(*Z*) (solid) and conditional mean function 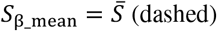. B. Scatterplot of overall performance *S*_ref_ against lateralization index *Z*_ref_ for reference data (Corr(*R*_ref_, *L*_ref_) ≅ 0), with LOESS regression line *S*_ref_loess_(*Z*) (solid), conditional mean function *S*_ref_mean_(*Z*) (dashed thick), and 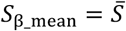 (thin dashed). C. The difference curves obtained from conditional mean functions *S*_β_mean_ − *S*_ref_mean_(*Z*). and from LOESS curves *S*_loess_(*Z*) − *S*_ref_loess_(*Z*). Regression lines in graphs A and B were calculated with the built in LOESS regression procedure (span = 0.6) of Mathcad Professional. Conditional mean function 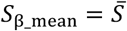 is a constant equal to the mean of performance (we assumed *S* and *Z* independence). The values of conditional mean function *S*_ref_mean_(*Z*) = *S*_mean_(*Z*; *ρ* = 0) for reference data presented in graph B were calculated from Equation 14.

## 3 Discussion and Conclusions

Leask and Crow (1997) asked a very important question: What is the true influence of laterality on performance, not obscured by the confounding effects of the mathematical, functional interdependence between the measures of the two? Having discussed the problems posed by the mathematical interdependence of the measures (independently described by Mazoyer and Tzourio-Mazoyer, 2004, recognized also by Chiarello et al., 2009), they devised a method (LCM) intended to remove the part of the dependence that is due to the functional, mathematical relation. Unfortunately, as we have shown above, the method produces virtually the same results for true observations as for simulated, random data. Particularly, it shows an apparent “laterality optimum” for simulated data where no causal relation between the indices exists, hence obviously no true optimum is possible. For random data with negative *R*, *L* correlation, the LCM reveals a nonexistent u-shaped relationship. In both simulations, the method produced curves very similar to the ones it provided for empirical data. The outcome of the LCM seems then to be mostly a mathematically predictable, artifactual appearance, born from subtracting the diverse nonlinear dependencies of the measures of laterality and performance. The mathematical relation between the measures changes with the size and the direction of the *R* and *L* correlation. Therefore, subtracting – as in the LCM algorithm – two curves derived from datasets that vary in this correlation results in a difference curve that reflects mostly the intricate complexities of this mathematical relation and cannot be rightly interpreted as depicting a true influence of laterality on performance.

Two factors are jointly responsible for the artifact in the Leask and Crow method: (1) the choice of the nonlinear function of the left- and right-hand performance measures for the independent variable; and (2) the use of the uncorrelated data to produce the reference against which local averages of the outcome variable are compared. It might seem at first that eliminating these factors, e.g. by using the unweighted *R* − *L* difference and a correlated reference, would repair the LCM method, but we do not know for sure. To assess the net asymmetry–performance relation by comparing the observed values with some “null” reference data, we need to specify the underlying null joint probability distribution (and parameters). Without a viable model of the lateralized processes that generate this distribution, what the divergence of the empirical data from the null reference reflects cannot be determined: it might be the sought-after net interaction of asymmetry and performance, the null model’s mismatch, sampling error, or a mixture of these^6^. Yet, considering that even something as elementary as the inter-trial variability of simple response times is mostly a mystery (Saron et al., 2003), we are not too optimistic about the prospects of obtaining a model of performance asymmetries that is detailed enough to match the differential logic of the LCM.

Among many other things, the missing model of laterality must define the direction of the expected relation between laterality and performance. The former is often believed to reflect a functional and structural brain asymmetry that has been selected in evolution due to the processing advantage it offers (e.g. Corballis, 2017). So then, laterality is presupposed to be the independent variable that, more or less directly, affects the dependent variable, performance. In the context of the issue of the adaptive value of asymmetry, such a notion seems quite natural. However, modern laterality research has questioned many if not most of the time-honored, ostensibly evident, assumptions: global asymmetry of the human brain structure has been shown to be unrelated to handedness (Good et al., 2001; Kong et al., 2018); large sample human study has shown handedness to be explained mostly by unique environmental effects (Vuoksimaa, Koskenvuo, Rose, & Kaprio, 2009); animal research suggests that population level laterality can be modified more by environmental than genetic factors (Giljov, Karenina, Ingram, & Malashichev, 2015; Giljov, Karenina, & Malashichev, 2012); many results undermine the notion of singular laterality by demonstrating the independence of various brain and functional asymmetries (Liu, Stufflebeam, Sepulcre, Hedden, & Buckner, 2009; Badzakova-Trajkov, Corballis, & Häberling, 2016; Häberling & Corballis, 2016; Mellet et al., 2014). In an important paper, Bishop (2013) argued that although atypical laterality had been traditionally believed to be responsible for language impairment in children, many other causal scenarios might explain the observed covariance, and that the most counter-intuitive of them, namely assuming laterality to be a consequence rather than cause of language impairment, is paradoxically quite a viable possibility. While a discussion on the concept of laterality and possible causes of the expected laterality–performance relation is certainly needed, we feel that, for now, an exploratory approach that does not presuppose any specific causal chain might be preferable.

Leask (2003) made a similar proposition. He explored the asymmetry of performance simply by plotting left- and right-hand performances against each other and approximating their relation by fitting a locally averaged principal component curve (he chose it over LOESS to avoid assumptions regarding which variable was dependent and which was independent). The resulting curve, fitted to the scatterplot of NCDS box-marking data, showed that the absolute difference between right- and left-hand performance rose linearly with overall performance, up to a point around the average, after which the difference remained relatively constant in spite of the increase in overall performance. Leask’s proposition of treating laterality as a bivariate concept with no assumptions regarding the direction of the dependence is very interesting. Its simplicity helps in obtaining an objective, assumption-free description of the laterality–performance relation. Leask’s analysis also avoids the potentially troublesome Laterality Index. The index’s formula with the *R* + *L* sum in the denominator – as popularized by its use in the Oldfield (1971) laterality questionnaire – has been adopted to compensate for the restricted variability of simple difference at the extremes of performance or preference (Boles et al., 2008; Bryden, 2012). In the 1970s, several authors debated the issue of laterality scaling and suggested that the ideal measure should be independent of performance (Birkett and Wilson, 1979; Colbourn, 1978; Marshall, Caplan, & Holmes, 1975)^7^. While in some applications such a measure might indeed be useful, analysis of the relation between laterality and performance should probably avoid a laterality measure designed specifically to be performance independent.

Abandoning the notion of laterality as a predictor of performance also encourages a less obvious way of imaging it. As performance is responsible for much more of the data dispersion visible on the *R*, *L* scatterplot than lateral differences^8^, it seems preferable to present the relation of the two, assigning the former to the x-axis, and the latter to the y-axis. Figure 4 presents plots of NCDS box-marking performance asymmetries expressed by *R* − *L* differences (on the left) and Laterality Index values (on the right) as functions of overall performance represented by *R* + *L* sums. The data were sorted by performance values and the dependences were approximated with LOESS. The *R* − *L* differences rise monotonically with performance, which is to be expected due to the mathematical relatedness of the measures. However, a less expected tendency for the rise to slow down above the median can also be seen. Similar nonlinearity is present for the Laterality Index, although due to the different scaling of the measure, here the curve is falling. This interesting effect is consistent with the observations of Leask (2003), and while it definitely seems intriguing, and there is a chance that it reflects something more than just a mathematical glitch, we cannot be certain without a feasible model of laterality. The model has to guide the analysis and dictate the choice of the asymmetry measure if the advantage or disadvantage of asymmetry is to be inferred from data like NCDS hand skill scores, where both performance and asymmetry measures are derived from the same observations^9^. We do not think that such a model is currently feasible, therefore we advocate cautious interpretation. Nevertheless, at least two conclusions seem justified by our examples and those of Leask (2003). Firstly, given the large dispersion of observations, as is evident in Fig. 4, the local approximations of the means do not seem to suggest any optimal laterality range for either laterality measure. This concurs with our earlier conclusion that the purported “optimum of laterality” (Leask and Crow, 1997; 2006) is mostly an artifact (as is the “advantage” proportional to absolute difference, described by Hirnstein et al., 2014). Secondly and perhaps more importantly, whatever the nature of the discussed association, it is very weak – most likely too weak to have much adaptive relevance.

**Figure 4.**
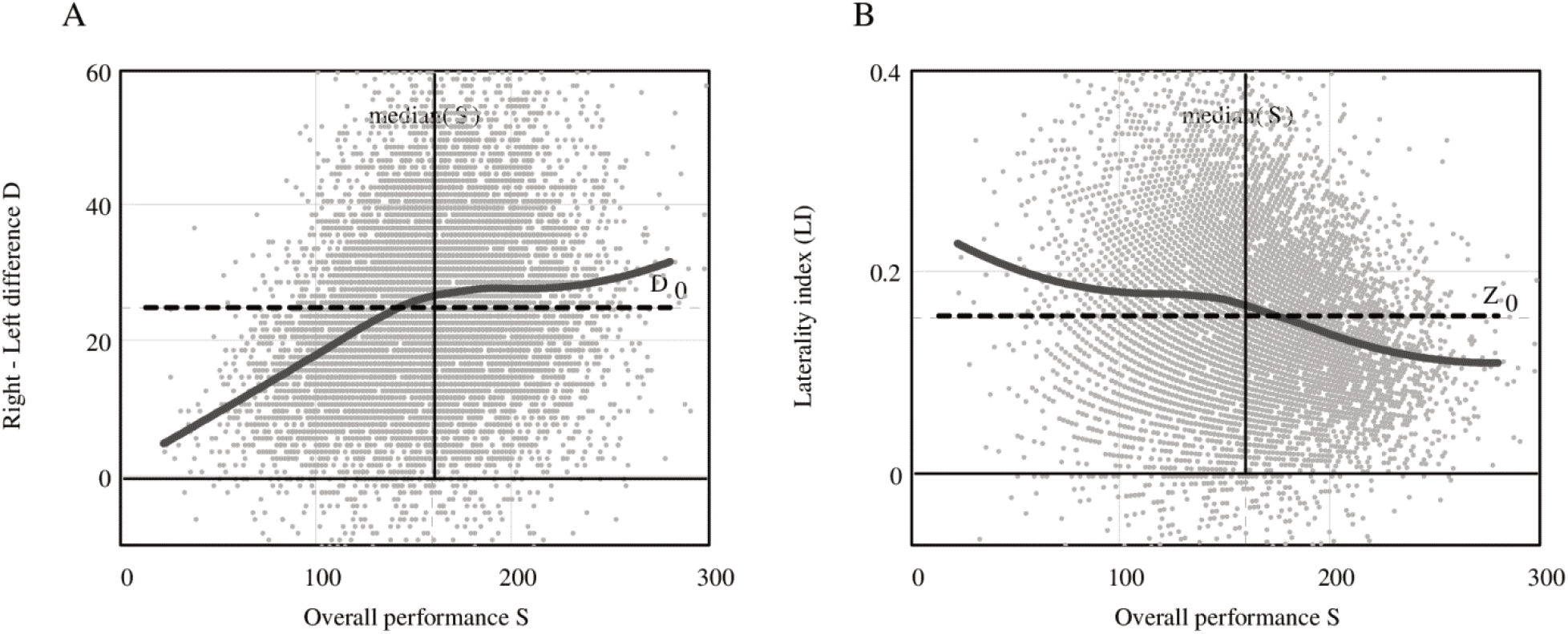
How does laterality change with performance? The scatterplots (grey dots) present *R* − *L* differences (graph A) and laterality indexes (graph B) for NCDS box-marking results (*N* = 11206; righthanders) as a function of increasing overall performance. Thick solid lines present local LOESS regression approximations of these relationships. The horizontal dashed lines and vertical thin solid lines present mean laterality and median performance levels, respectively.

## 4. Appendix

The aim of the present work was to determine if the LCM algorithm applied to random data produced effects similar to those reported for true observations. We chose binormal distribution for the simulated hand skill, and beta distribution for the simulated dichotic listening scores. These distributions were chosen due to their reasonable resemblance to the measured data and their relative algebraic simplicity. We should put a caveat here that since no theoretical model of laterality guided this decision, it would not be appropriate to treat the random dataset normatively and assign any theoretical meaning to the departures of the real data from it.

## 4.1 Distribution of box-marking scores – bivariate normal

As mentioned above, when modeling the observed left- and right-hand NCDS box-marking performances with two correlated random variables *R*, *L*, we chose a bivariate normal distribution for the generated “scores”. For positive values *x*_R_, *x*_L_ of *R*, *L* we used the joint probability density function of the bivariate normal distribution in the form:

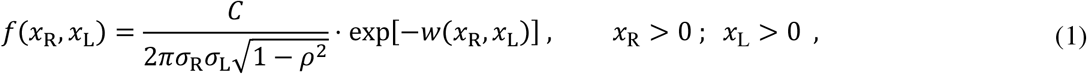

where the argument of the exponential function was defined as:

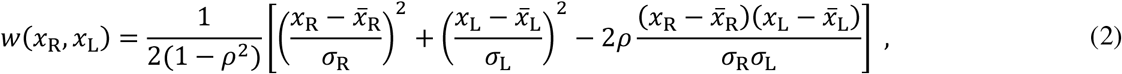

and the constant *C* was determined by normalization condition 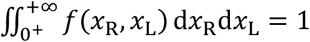 In the formulae given in Equations 1 and 2, the overlined symbols 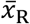 and 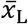 denote means of *x*_R_ and *x*_L_, *σ*_R_ and *σ*_L_ denote their standard deviations, and *p* denotes the coefficient of linear correlation between them. An approximation using the binormal probability density function only for positive values of variables is acceptable if the means of the variables in question are much greater than their standard deviations, as is the case with the NCDS box-marking data (where 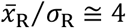, 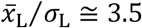; see Table 1). While in general, a zero value of the linear correlation coefficient does not necessarily imply independence, in cases like ours, where *x*_R_ and *x*_L_ are subject to a bivariate normal distribution, their independence does follow from the correlation *ρ* = 0.

**Table 1.** Estimates of means 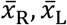, standard deviations *σ*_R_, *σ*_L_ and correlation coefficient *ρ* for randomly generated data (N = 12 000) with and without correlation, compared to the estimates for the NCDS box-marking results.

| Parameter | Estimate derived from the NCDS observations | Descriptive statistics for the generated datasets: |  |
| --- | --- | --- | --- |
|  |  | correlated | uncorrelated |
| $\bar{x}_R$ | 92.0 | 91.9 | 92.3 |
| $\sigma_R$ | 22.6 | 22.4 | 22.7 |
| $\bar{x}_L$ | 66.9 | 67.2 | 67.0 |
| $\sigma_L$ | 19.4 | 19.3 | 19.2 |
| $\rho$ | 0.782 | 0.778 | 0.005 |

## 4.2 Random dataset generation

To obtain a set of correlated random variable pairs (*R*_g_, *L*_g_) simulating right- and left-hand NCDS box-marking “scores”, we used Mathcad Professional’s built-in functions to independently generate normally distributed variables *y*_A_~N(0,1) and *y*_B_~N(0,1). Then, we linearly transformed the generated variables using the formulae from Walck (2007):

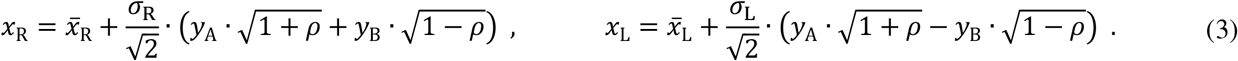

The formulae ensured that the calculated variables *x*_R_ and *x*_L_ had the bivariate normal distribution defined in Equation 1. For numerical calculations based on Equation 3 we used estimates for mean values 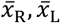, variances 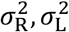 and correlation coefficient *p* matching the actual NCDS box-marking scores. Analogically, we generated a set of uncorrelated (*ρ* = 0) data, (*R*_o_, *L*_o_). The two datasets are shown as scatterplots in Figure 5, while the distribution parameters of the variables *x*_R_ and *x*_L_ are listed in Table 1.

**Figure 5.**
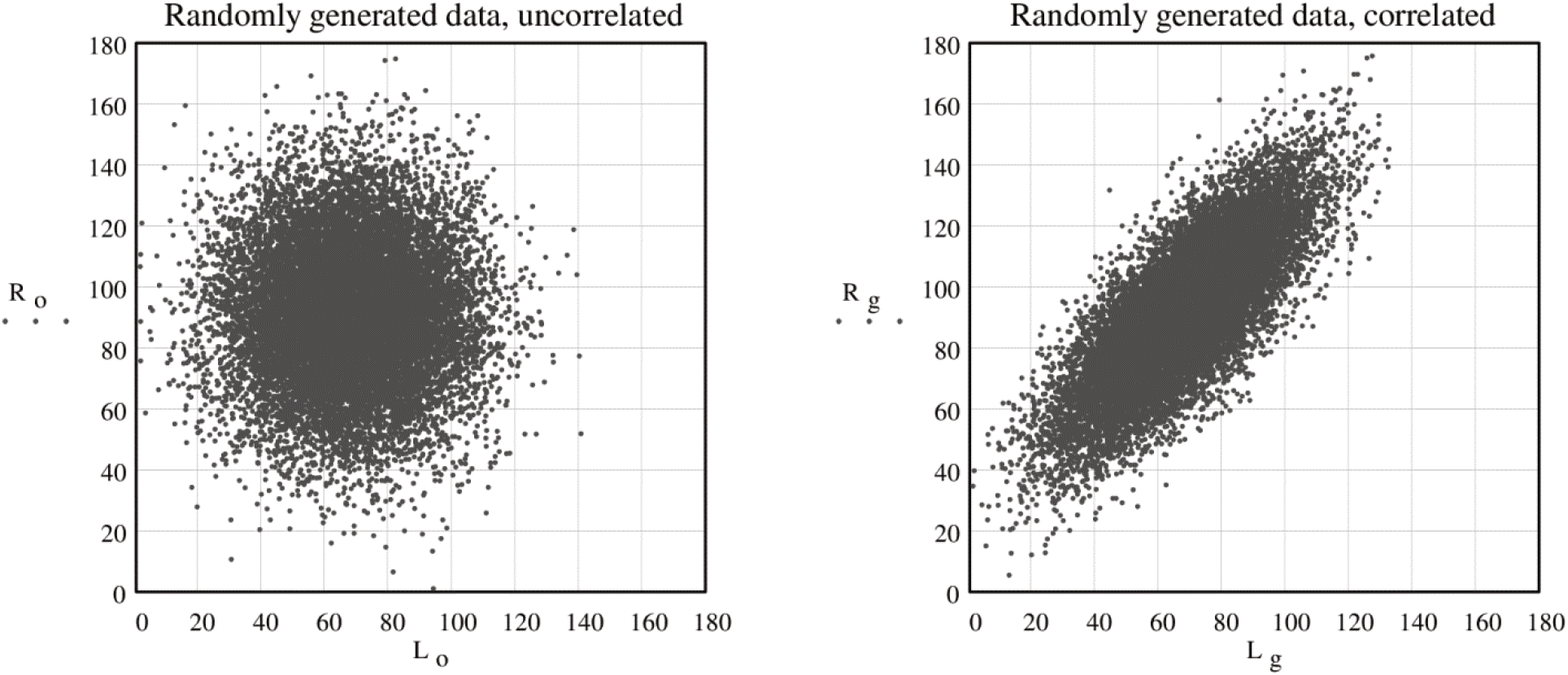
Scatterplots of two datasets for two randomly generated variables *R* = *x*_R_, *L* = *x*_L_ (right-s and left-hand scores), plotted against each other. Both visualized datasets were generated using the transformation shown in Equation 3, both with the same distribution parameters as given in the NCDS box-marking data (see Table 1), but the left and right scatterplots show uncorrelated (*ρ* = 0) and correlated (*ρ* = 0.782) variables, respectively (built-in Mathcad Professional software procedures were used).

## 4.3 Conditional probability density function for overall performance for given value of laterality index

The LOESS regression used in the LCM to approximate the relation between laterality and performance can be seen as an estimate of the conditional mean (or also mode in the case of symmetrical distributions) function. Therefore, the effects revealed by regression can also be predicted analytically. To that end we derived the conditional mean and conditional mode formulae for *R* and *L* using the conditional probability density function for overall performance given fixed values of laterality index (LI).

Although it is possible to formulate the conditional probability density function for variables *x*_R_, *x*_L_ for given values of LI, it is easier to determine this density function of their sum *S* (overall performance) in order to specify a conditional mean function of *S*. Then one can determine the conditional mean functions for the initial measures, i.e. performance of the right and left hand, keeping in mind that the relationship between *x*_R_, *x*_L_ and their sum *S* is linear. An analogical method can be used to determine the conditional mode functions.

We needed to specify the joint probability density function for the new variables, overall performance *S* and laterality index (LI, for which we choose symbol *Z*), defined by one-to-one transformation of the original variables:

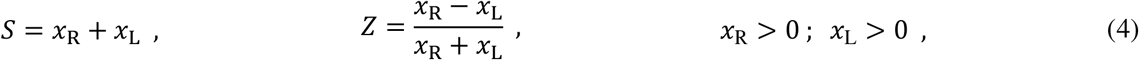

for which the inverse transformation to the original variables is

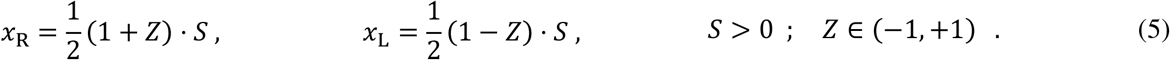

The following estimators of mean values of the new variables *S*, *Z* and variance 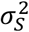 of variable *S* were used:

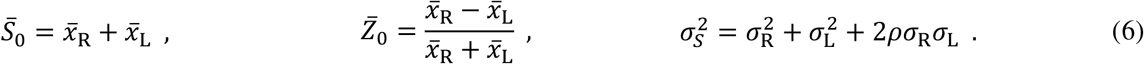

Moreover for more transparency and clarity of the formulae (which will be derived later), the two dimensionless parameters *ε*, *α* are defined as:

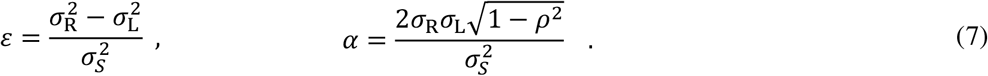

Then we were able to transform the original probability distribution of variables *x*_R_, *x*_L_ to the probability distribution of the new variables just defined in Equation 4: overall performance *S* and the laterality index *Z*. This required the derivation of a joint probability density function *g*(*S*, *Z*) for overall performance S and laterality index *Z*. The function must satisfy the following relation:

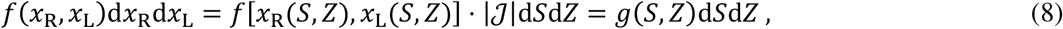

where 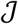 is the Jacobian for the inverse transformation Equation 5 and its absolute value is 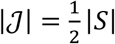 for *S* ≠ 0.

From Equations 1, 2 and 8, the joint probability density function for overall performance *S* and laterality index *Z* was derived as follows:

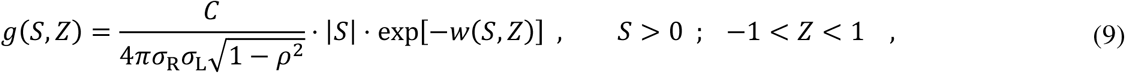

where *w*(*S*, *Z*) derived from Equation 2, and Equations 5–7 are given by the formula:

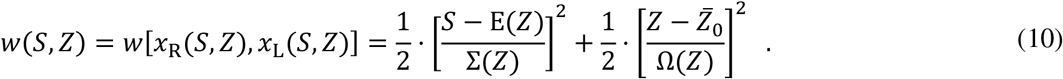

Functions E(*Z*), Σ(*Z*), Ω(*Z*) in Equation 10 are defined as:

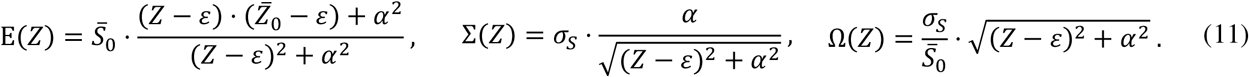

for which the following equalities are true: 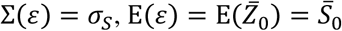 and 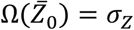 (i.e. standard deviation of *Z*).

The conditional probability density function of *S* (sum, overall performance) for fixed values of the second variable *Z* (laterality index LI), normalized with respect to the first one *S*, is denoted by *g*_*s*_(*S*|*Z*) for given values of *Z*. This function is defined by the following relation:

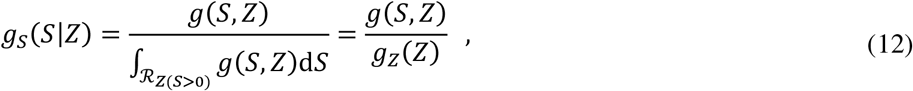

which expresses Bayes’ theorem *g*_*s*_(*S*|*Z*) · *g*_*z*_(*Z*) = *g*(*S*, *Z*). Function *g*_*z*_(*Z*) in Equation 12 is the so-called marginal distribution defined as integral 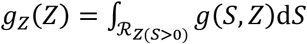, where 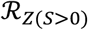 denotes the set of all points in the range 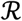 of (*S*, *Z*) for a given value of *Z* (in each region *S* > 0 and *Z* ∈ (−1, +1)).

## 4.3.1 Conditional mean function

To derive the conditional mean functions of the left- and right-hand performances, we used the conditional mean function for overall performance *S*_mean_(*Z*), which in turn was determined according to the definition of the mean value of *S* when using the conditional probability density function *g*_*s*_(*S*|*Z*) of overall performance *S* for fixed values of the second variable *Z* (laterality index LI):

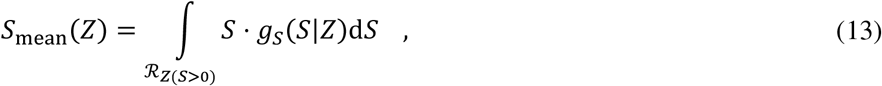

where the conditional probability density function *g*_*s*_(*S*|*Z*) is defined by Equation 12. The resulting formula for the conditional mean value of overall performance *S*_mean_(*Z*) takes the form:

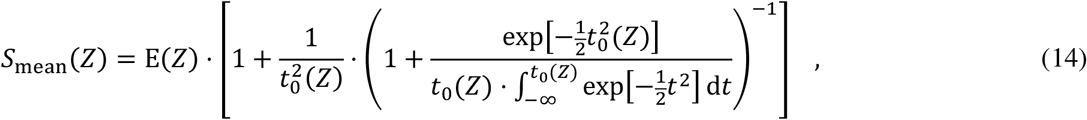

where E(*Z*), Σ(*Z*) denote functions defined earlier in Equation 11 and *t*_0_(*Z*) = E(*Z*)/Σ(*Z*).

Conditional means *x*_R_mean_(*Z*), *x*_L_mean_(*Z*) for right- and left-hand performance, being the functions of laterality index *Z* on the basis of Equation 5, can be written as follows:

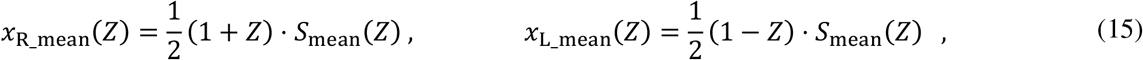

where the conditional mean function for overall performance *S*_mean_(*Z*) is defined by Equation 14. The above derivation of the conditional mean functions on the basis of Equation 5 was possible because the overall performance *S* is a linear combination of the variables *x*_R_, *x*_L_, i.e. their sum (see Equation 4). The formula for *S*_mean_(*Z*) given in Equation 14 is not a closed-form expression because it involves an integral which for each value of *Z* must be either computed numerically or otherwise approximated. If the variables *x*_R_, *x*_L_, *S* underlie symmetrical probability distributions, it is possible to replace the conditional mean function with the conditional mode function, which has nearly the same trajectory as the former, but has a formula that makes calculations simpler.

## 4.3.2 Conditional mode function

Conditional mode function for overall performance *S* given *Z* value (laterality index LI) may be defined as follows: *S*_mode_(*Z*) = arg max_*s*∈*R*_ *g*_*s*_(*S*|*Z*); therefore, it can be determined by finding for which value *S* = *S*_mode_(*Z*) (conditional mode); the conditional density function *g*_*s*_(*S*|*Z*) has a maximum for a given value of *Z*. As a result we obtain the function for the conditional mode of overall performance:

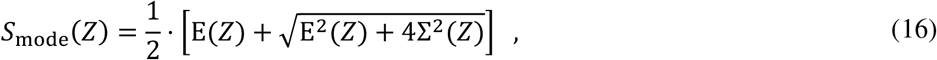

where E(*Z*), Σ(*Z*) denote functions defined earlier in Equation 11. Conditional modes *x*_R_mode_(*Z*), *x*_L_mode_(*Z*) for right- and left-hand performance can then be expressed as the function of laterality index *Z*, in accordance with Equation 5:

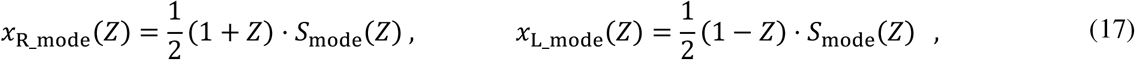

where the function *S*_mode_(*Z*) is defined by Equation 16.

In Figure 6 we compare the curves representing the derived functions for conditional means (Equation 15) and modes (Equation 17) with the locally weighted regression (LOESS) curve for a randomly generated dataset with distribution parameters matching the values observed in NCDS data. As expected for symmetric probability distributions, conditional mean and mode functions are nearly identical, and both are well approximated by the local regression LOESS line. In our own analyses we chose conditional mean over mode (cf. Section 2.1); however, the latter might also be used – and perhaps even preferred due to its computational simplicity – as the probability distributions *x*_R_, *x*_L_, *S* are symmetric.

**Figure 6.**
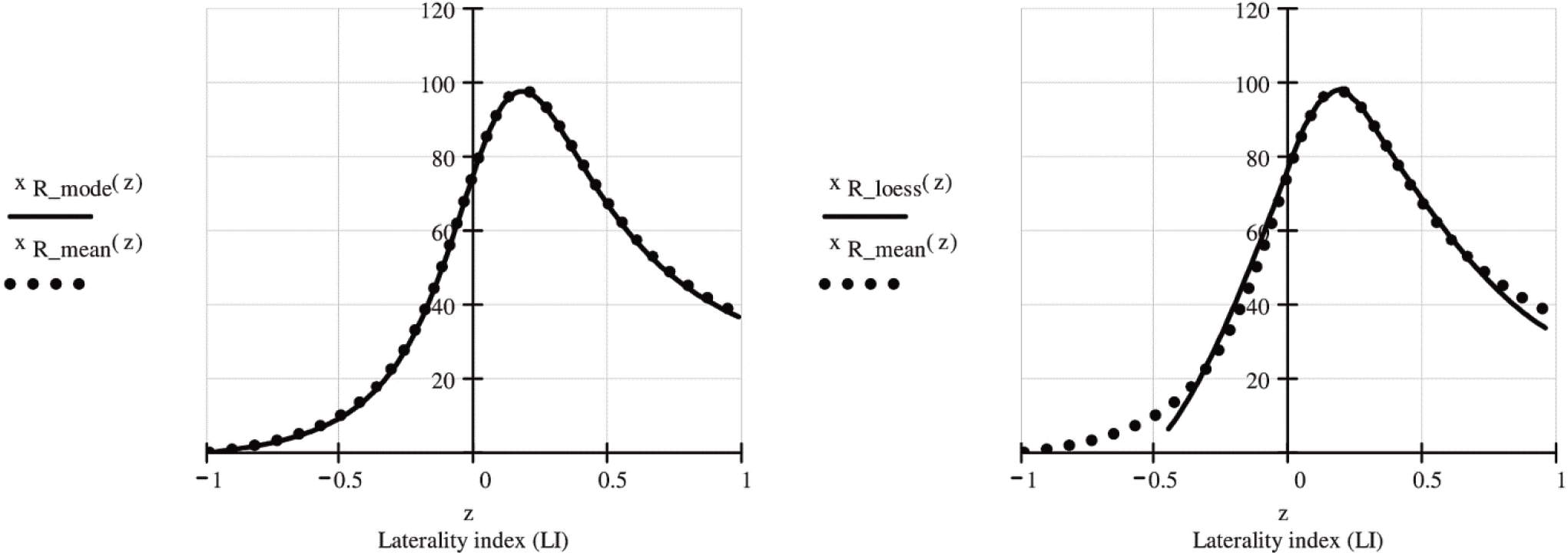
Graphical comparison of the conditional mean and mode functions and the LOESS regression line. Left: the conditional mode *x*_R_mode_(*Z*) (solid line) and conditional mean *x*_R_mean_(*Z*) (dotted line) functions are plotted against the lateralization index Z. Right: the courses of the LOESS regression line *x*_R_loess_(*Z*) (solid) and the conditional mean function *x*_R_mean_(*Z*) (dotted line) are compared. The presented functions were calculated on the basis of the formulae shown in Equations 15 and 17 with the same distribution parameters as given in the NCDS box-marking data (see Table 1), and the LOESS line was fitted with span 0.6 for randomly generated and correlated (*ρ* = 0.782) data (built-in Mathcad Professional software procedures were used).

## 4.4 Difference functions

Leask and Crow (1997; 2006) compared the values obtained with LOESS regression for correlated variables *x*_R_, *x*_L_ to reference data derived from uncorrelated variables. We performed similar comparisons for actual observations, for randomly generated data, and for conditional mean functions. To this end, we used difference functions defined as differences between the values of the performance functions for variables *x*_R_ or *S* with (*ρ* ≠ 0) and without (*ρ* = 0) correlation for the same value of lateralization index *Z*. We defined the difference function Δ*x*_R_mean_(*Z*) for the conditional means for right-hand performance *x*_R_ as:

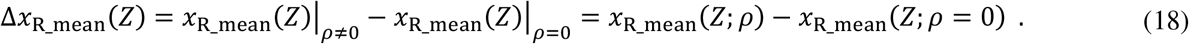

Analogous difference functions can be determined for conditional mode functions or values predicted by LOESS regression in randomly generated or empirical real data.

It is noteworthy that for randomly generated data with a specified joint probability density function (Equations 1, 2), the difference functions for conditional means or modes described by Equations 18 and 15 or 17 are strictly defined functions of the variable *Z* (laterality index). Moreover these functions are defined by the statistical parameters of the probability distribution: 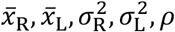. It follows that with numerical methods or mathematical analysis alone one can test these functions for the presence of maxima (“optima of laterality”) and obtain the laterality index values at which these maxima occur.

## 4.5 Dichotic Listening – beta distribution

Testing the LCM method with randomly generated data simulating the results of the dichotic listening experiment of Hirnstein et al. (2014), we modeled the scores using the following definitions of variables *R*, *L* (for right- and left-ear performance respectively):

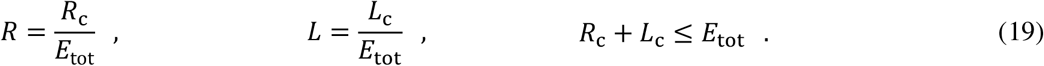

Additionally, variables *S*, which denotes overall performance, and *W*, which denotes relative right-ear efficiency, were calculated according to the formulae:

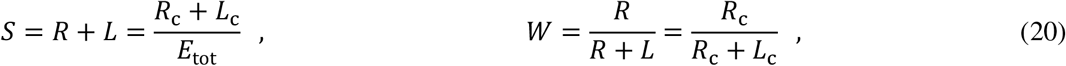

where *R*_c_, *L*_c_ stand for the numbers of correctly identified syllables for the right and left ear respectively, and *E*_tot_ = 30 is the total number of possible correct responses. It follows from these definitions that the values of variables *S*, *W* stay within the [0,1] interval. According to Hirnstein et al. (2014), the correlation between *S* and *W* was negligible, i.e. Corr(*S*, *W*) = Corr(*S*, *Z*) = 0.06 (where *W* relates to the lateralization index *Z* as follows: *Z* = 2 · *W* – 1). Therefore it seemed reasonable to assume that the variables S and W were independently subject to two separate beta distributions.

A standard beta distribution for variable *x* with the shape parameters *p* ∈ (1, ∞) and *q* ∈ (1, ∞) is a continuous distribution determined for *x* ∈ [0,1] with a probability density function given by the formula:

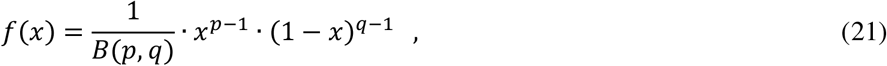

where the beta function *B*(*p*, *q*) is defined as 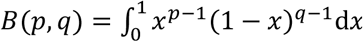. The estimation of the beta distribution parameters *p*, *q* is possible thanks to known estimates of mean value and standard deviation 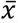, *σ*_*x*_ with the formulae:

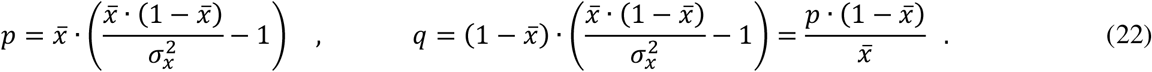

As only the distribution parameters for the *R*, *L* measures were available (i.e. the set of estimates 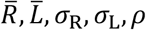), we estimated means and variances for variables *S*, *W* with the following:

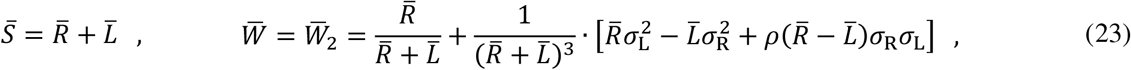

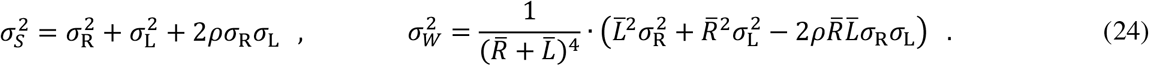

Substituting the data from Hirnstein et al.’s (2014) study (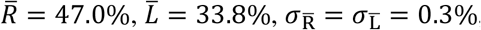, *N* = 1839, Corr(*R*, *L*) = *ρ* = −0.51, where 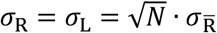) into Equations 23 and 24, we obtained standard deviations *σ*_s_ = 12.7%, *σ*_*w*_ = 13.9% and means 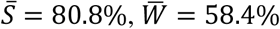. Using these values we determined (cf. Equation 22) the parameters of the two separate beta distributions, to which the random generated variables S and W were subject. We then used Mathcad professional to randomly generate data (*N* = 1839) subject to beta distributions: *S*~Beta(*p*_*s*_, *q*_*s*_), *W*~Beta(*p*_*w*_, *q*_*w*_). Values of the *R*, *L* and *Z* variables were calculated from:

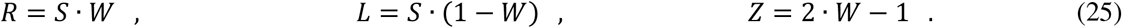

As result we get a dataset of correlated variables *R*, *L* with distribution parameters 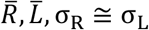 and correlation *ρ* = Corr(*R*, *L*) ≅ −0.5 that agree well with the empirical values. The outcome of this simulation (*N* = 1839) is presented in Figure 7 with histograms where the marginal probability density functions for the variables *R*, *L* and S are additionally plotted.

**Figure 7.**
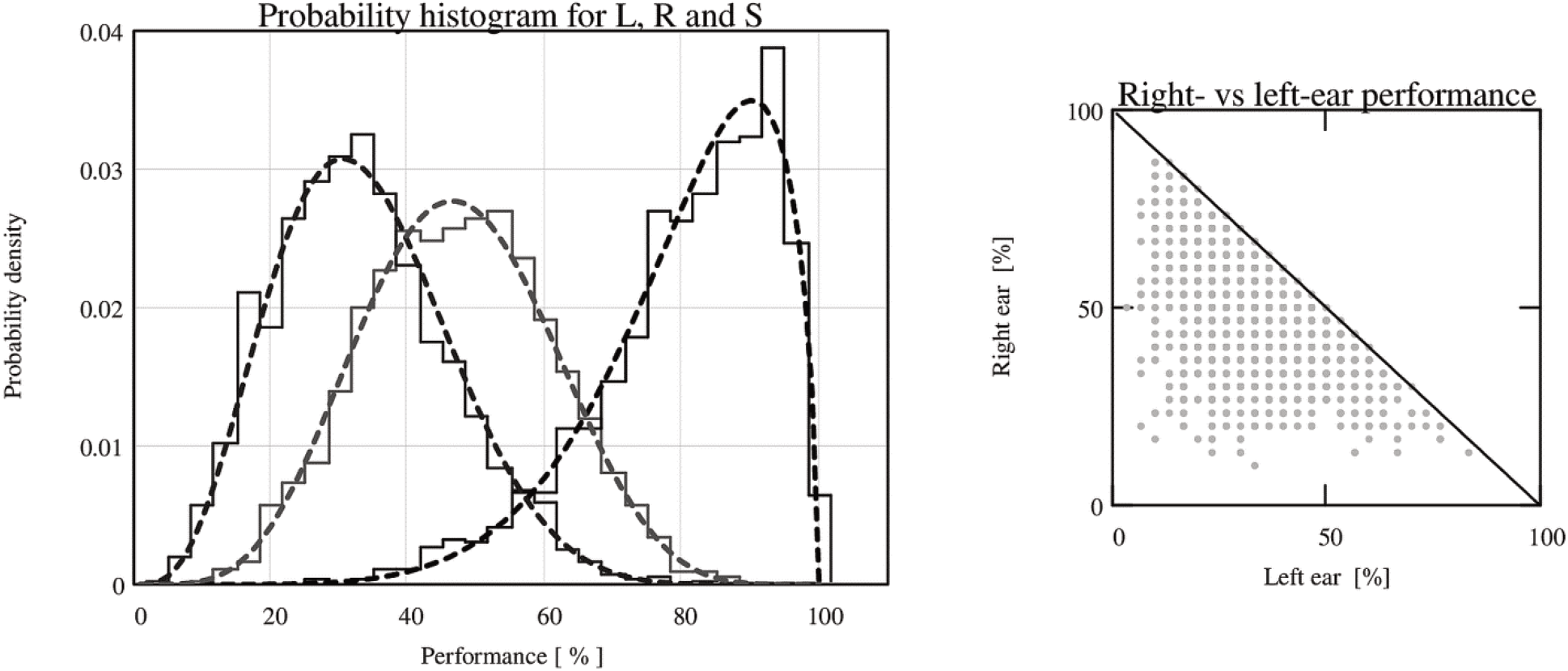
Simulation outcome of the dichotic listening experiment scores (non-forced condition). Left: histograms of probability density for generated variable S = *R* + *L* describing overall performance and for calculated variables *R*, *L* for right- and left-ear performance. Right: scatterplot of right-ear performance *R* plotted against left-ear *L* performance. Data points falling on the diagonal represent 100% overall performance *S*.

Noncorrelated reference data were created according to the decorrelation procedure applied by Hirnstein et al. (2014), where *R* and *L* scores are shifted with respect to each other. On the basis of the shifted score set *R*_ref_ = *R*, *L*_ref_ = *L*_(shifted)_, we computed reference data for overall performance *S*_ref_ and the lateralization index *Z*_ref_.

After decorrelation (Corr(*R*_ref_, *L*_ref_) ≅ 0.01), the overall performance *S*_ref_ = *R*_ref_ + *L*_ref_ was distributed approximately normally (see Figure 8). Thus the condition underlying the procedure for determining the conditional mean function (Equation 14) was satisfied. Therefore, for the analysis of reference data *S*_ref_, *Z*_ref_ we were able to reuse the already derived conditional mean function *S*_ref_mean_(*Z*) = *S*_mean_(*Z*; *ρ* = 0) by exploiting the formula defined in Equation (14) with zero correlation (*ρ* = 0) and distribution parameters for the parts defined in Equations 6 and 7, 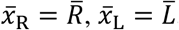, *σ*_R_ = *σ*_L_, as estimated for the analyzed data. For correlated data, the conditional mean function for overall performance *S* given *Z* value (laterality index LI) is a constant function 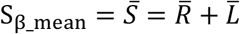 corresponding to the mean value of 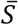 for independent random generated data *S* and *Z* = 2 · *W* − 1 giving the correlation Corr(*R*, *L*) ≅ −0.5 between *R* and *L* variables.

**Figure 8.**
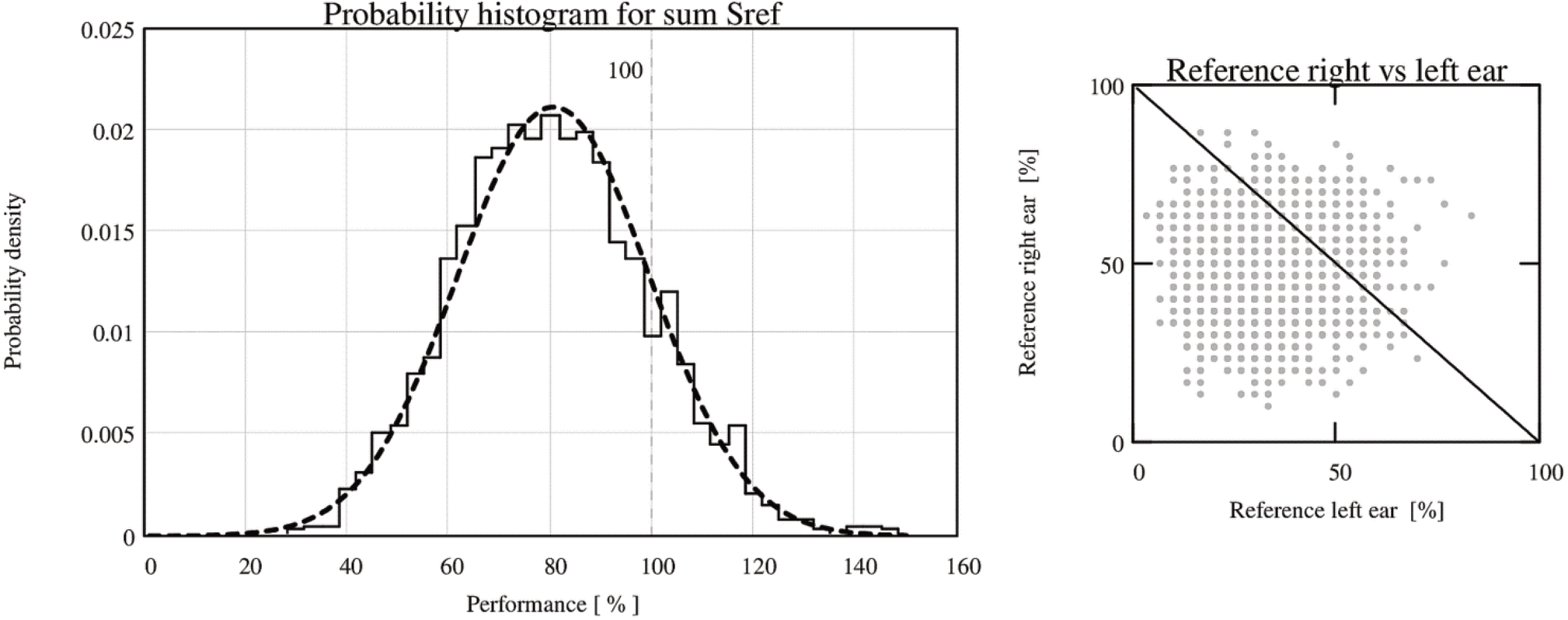
Uncorrelated reference data used in the simulation of the LCM algorithm for the dichotic listening results. Left: probability density histogram and function for reference variable *S*_ref_ = *R*_ref_ + *L*_ref_ describing overall performance. Right: scatterplot of right- and left-ear performances *R*_ref_ = *R*, *L*_ref_ = *L*_(shifted)_ plotted against each other. Data points falling on the diagonal represent the performance *S*_ref_ value of 100%. Notice that all points above the diagonal represent results that are impossible in real measurements.

## Footnotes

2 Box-marking scores for right-handed subjects from *National Child Development Study Response and Outcomes Dataset, 1958– 2013* [http://dx.doi.org/10.5255/UKDA-SN-5560-3] openly accessible at UK Data Archive

3 We generated binormally distributed random values shown in Figure 5 using the formula described in Equation 3. From the simulated right *R*_g_ and left *L*_g_ hand performance scores, the Laterality Index values *Z*_g_ = (*R*_g_ − *L*_g_)/(*R*_g_ + *L*_g_) seen on the scatterplot in Figure 1B were computed.

* No part of the study procedures or analyses was pre-registered in an institutional registry prior to the research being conducted

4 Minor differences in shapes of the curves computed for real (Figure 1A) and random (Figure 1B, C) data – similar to those actually present – are expected, considering the imperfect match between the empirical distribution of the observed *R* and *L* performance scores and the theoretical bivariate normal distribution used for random generation of correlated test data and derivation of conditional mean functions (cf. *Appendix*). It should also be mentioned that any quantitative comparisons of the curves will be confounded by the strong non-linear dependency between the lateralization coefficient LI and the observed values of *R* and *L*. For these reasons we think that only qualitative comparison is adequate here.

5 For compatibility with Hirnstein et al. (2014) we use sum *S* = *R* + *L* rather than just *R* as performance measure, and consequently we also use conditional mean functions for overall performance.

6 Of course the logic is different when one does what we did in the present work, i.e. it tests the method that has already been proposed. In this case, the striking similarity of the outcomes for real and for random data is alarming regardless of the distribution.

7 Despite broad adoption of the Laterality Index, the consensus has never been universal. Notably, Marian Annett, a major laterality expert, wrote in her monograph: “In my view, ‘indices’ and ‘quotients’ of hand preference put a gloss on ignorance. They are pseudoscience because they assign numbers without objective referents and therefore obscure what they purport to describe” (Annett, 2002, p. 26).

8 Pearson correlation of performances sum S with the first principal component derived with PCA, is close to 1 (for NCDS data *r* = 0.99964).

9 In a similar vein, McManus, Van Horn and Bryden (2016) explored the relation between hand skill asymmetry and performance using different tasks and laterality indices. In their interesting paper they discuss reasons for the intriguing invariance of the R–L difference relative to task difficulty and performance they found in the Tapley and Bryden (1985) task. This invariance is restricted to asymmetry expressed by unweighted R–L difference but not by LI. McManus et al. provide an extensive discussion which clearly shows what we also try to convey here: the relevance of the theory to the interpretation of the asymmetry–performance relation and the choice of the asymmetry measure.

Disclosure statement The authors have no conflicts of interest to disclose

